# Optogenetic control of cell morphogenesis on protein micropatterns

**DOI:** 10.1101/563353

**Authors:** Katja Zieske, R Dyche Mullins

## Abstract

Cell morphogenesis is critical for embryonic development, tissue formation, and wound healing. Our ability to manipulate endogenous mechanisms to control cell shape, however, remains limited. Here we combined surface micropatterning of adhesion molecules with optogenetic activation of intracellular signaling pathways to control the nature and morphology of cellular protrusions. We employed geometry-dependent pre-organization of cytoskeletal structures together with acute activation of signaling pathways that control actin assembly to create a tool capable of generating membrane protrusions at defined cellular locations. Further, we find that the size of microfabricated patterns of adhesion molecules influences the molecular mechanism of cell protrusion: larger patterns enable cells to create actin-filled lamellipodia while smaller patterns promote formation of spherical blebs. Optogenetic perturbation of signaling pathways in these cells changes the size of blebs and convert them into lamellipodia. Our results demonstrate how the coordinated manipulation of adhesion geometry and cytoskeletal dynamics can be used to control membrane protrusion and cell morphogenesis.

---

The ability of animal cells to dynamically control their shape underlies many essential processes, including embryonic development, tissue morphogenesis, and amoeboid cell migration. A key element in controlling cell shape is the ability to form membrane protrusions of defined morphology at specific locations. Dynamic membrane protrusions enable animal cells to interact with their environment in various ways: probing local mechanical properties, communicating with neighboring cells, and navigating complex three-dimensional matrices. Cellular protrusions themselves display distinct shapes, such as linear filopodia, sheet-like lamellipodia, or spherical blebs. These structures also rely on different molecular mechanisms.(1)(2) Filopodia and lamellipodia, for example, are created by the assembly of actin filaments under the control of localized cellular signals. Bleb formation, in contrast, is driven by hydrostatic pressure and is sensitive to the strength of the connection between the plasma membrane and the underlying cortex. Although individual protrusion morphologies have been extensively studied and many of the underlying signaling proteins are known (1)(3), we are just starting to understand mechanisms for transitions and molecular relationships between distinct protrusions.

To form a functional tissue animal cells must both create and make sense of complex, three-dimensional environments. Recent studies demonstrate that the geometry of molecular adhesion and physical confinement modulate the nature and morphology of membrane protrusions, especially those associated with cell motility.(4) These results help us understand how physical aspects of the external environment influence cell morphology, but they also raise many additional questions. We know very little, for example, about how environmental geometry affects morphology *transitions.* In addition, although properties of the cellular microenvironment can influence morphology, certain types of protrusion have been described as cell-type specific. One example is the observation of large, travelling blebs in some cell types(5)(6), which are morphologically distinct from the small, non-propagating blebs found in other cells. Theoretical concepts provide insight into how bleb size could be modulated by physical parameters, such as cortex contractility or nucleation size of blebs (7), but how do cells control these parameters to transition between protrusion morphologies? Despite decades of documenting cellular blebs and identification of key molecules for bleb formation(8), it is still unclear whether and how cells regulate bleb size. Is bleb size determined by distinct mechanisms and protein composition or does bleb size represent different states of the same mechanism? And how do signals from the extracellular environment contribute to this process?

To understand cell morphogenesis from a systems level new approaches aiming at uncovering fundamental parameters for cell shape determination and gaining custom control of cell morphologies are required. Protein micropatterning represents a powerful tool to mimic the extracellular environment and study how geometry and adhesion influences spatial organization of cellular components(9). Optogenetic methods for perturbing protein activity give us spatio-temporal control of intracellular signals, and have been used, for example, to induce light-dependent formation of lamellipodia(10). However, in the context of cell morphogenesis the synergistic potential of combining these techniques has not yet been explored or exploited.

Here, we combined optogenetics and engineered protein micropatterns to gain new levels of control over the mechanism and morphology of membrane protrusions. Integrating these two techniques allows to induce distinct protrusion morphologies, including small blebs, large blebs and lamellipodia. We demonstrate that this approach provides *temporal* control over protrusion morphology as well as control over the number and location of light-induced protrusion. In addition, we applied the combination of optogenetics and micropatterns to determine conditions for morphology transition. Specifically, we demonstrate an increase of blebbing activity with decreasing protein pattern size and transitions of protrusion morphology towards larger blebs or lamellipodia upon light-activation of signaling pathways.

Our results demonstrate a new level of engineering control over cellular protrusions and reveal that the geometry of cell adhesion and intracellular protein activation provide a parameter space in which diverse morphologies of membrane protrusions can be generated.

## Heterogeneity of light-induced membrane protrusions along cell boundaries

To explore the synergistic regulation of cell morphology by microenvironments and intracellular signals we first expressed a photactivatable signaling molecule capable of inducing actin-based protrusions in cells on homogeneous substrates. Briefly, we stably transfected HFF1 cells, which form distinctive lamellipodia, with mCherry-tagged, photoactivatable Rac1 (PA-Rac1), a protein previously found to induce membrane protrusions at locations exposed to 458- or 473-nm light(10) (Figure 1A). We compared cell spreading under control conditions (Figure 1B) and light-induced activation of PA-Rac1 (Figure 1C). We accessed cell expansion by subtracting the PA-Rac1 signal at the starting timepoint of the experiment from the signal after 2 or 5 minutes. We observed little cell expansion under control conditions, but significant cell expansion when cells were uniformly activated with blue light. Interestingly, spreading was not uniform around the cell boundaries but exhibited significant degrees of spatial heterogeneity, which classify into three morphologies: (i) large (~5 μm) lamellipodia, (ii) regions with moderate lamellipodia protrusion (<5 μm), and (iii) quiescent regions with no detectable lamellipodial protrusions (Fig, 1Ci-iii).

**Fig 1:**
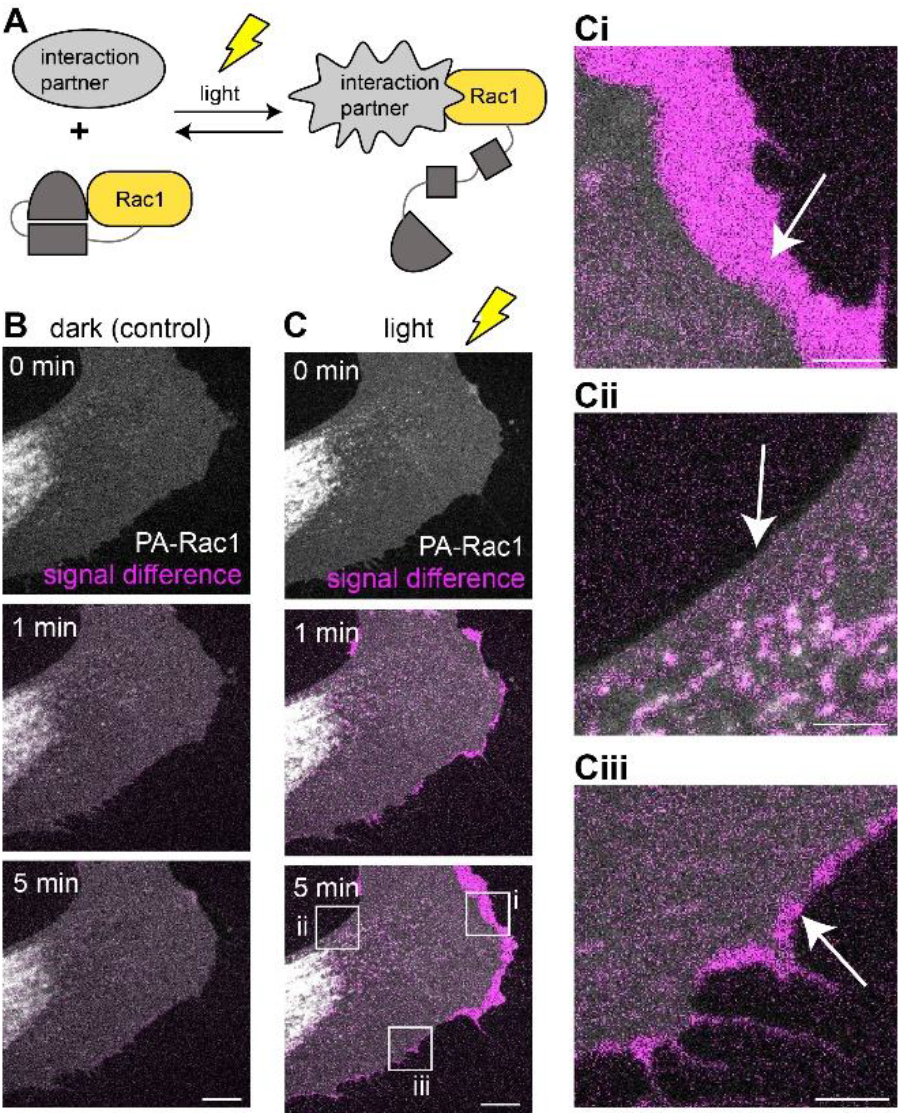
Homogeneous optogenetic activation of Rac1 results in heterogeneous formation of cell protrusions. (A) Photoactivatable Rac1 fused to n-terminal mCherry (PA-Rac1)(10) was stably expressed in HHF1 cells using lentivirus transfection. (B,C) Confocal time-lapse images of PA-Rac1 distribution (white) show cell areas and their temporal development under dark (B) and photoactivating conditions (C). As a measure for cellular protrusion formation the signal difference of PA-Rac1 at later time points (1min, 5 min) and PA-Rac1 at the beginning of time-lapse acquisition (0 min) was computed (magenta). White boxes indicate locations of magnified regions shown in (Ci-iii). Note that protrusion formation is heterogeneous along the cell boundary. (B,C) Scale bar: 20 μm, (Ci-iii) Scale bar: 5 μm

These fluctuations of lamellipodia expansion along the cell boundary demonstrate that the spatial efficiency of optogenetically-driven cell functions can be heterogeneous, despite homogeneous illumination. A likely reason for this observation are the heterogeneous biochemistry and cell mechanics along cellular boundaries. Notably, however this heterogeneity of protrusion formation upon optogenetic activation could provide exiting new opportunities for controlling cell protrusions. Specifically, the number of protrusion and their location may be controllable by employing whole cell illumination. For this goal however, engineering controlls need to be found to spatially control the source of optogenetic heterogeneity.

### The cellular microenvironment modulates responses to optogenetic activation of signaling molecules

One explanation for the spatial heterogeneity of protrusions formed in response to whole-cell optogenetic activation of Rac1 could be the heterogeneous distribution of pre-existing cytoskeletal networks in the target cell. Membrane-proximal actin bundles (stress fibers), for example, could locally blunt protrusive responses(11) whereas cortical networks with more randomly organized filaments could be permissive for protrusion. This non-genetic variability might be considered a form of ‘cytoskeletal epigenetics’(12). To test this hypothesis, we exploited a relationship between adhesion geometry and actin filament organization to achieve spatial control over the cytoskeleton near the plasma membrane. Thery et al.(11) previously showed that fibroblasts plated on triangular micro-patterns of adhesion molecules display characteristic actin distributions, with thick bundles along the sides and dynamic branched networks at the corners(11). Therefore, we imaged protrusion formation in response to global optogenetic activation of Rac1 in fibroblasts plated on micro-patterned triangles, generated using laser lithography. In these experiments the constrained adhesion geometry caused preferential formation of membrane protrusions at cell corners compared to sides (Fig 2A).

**Fig 2:**
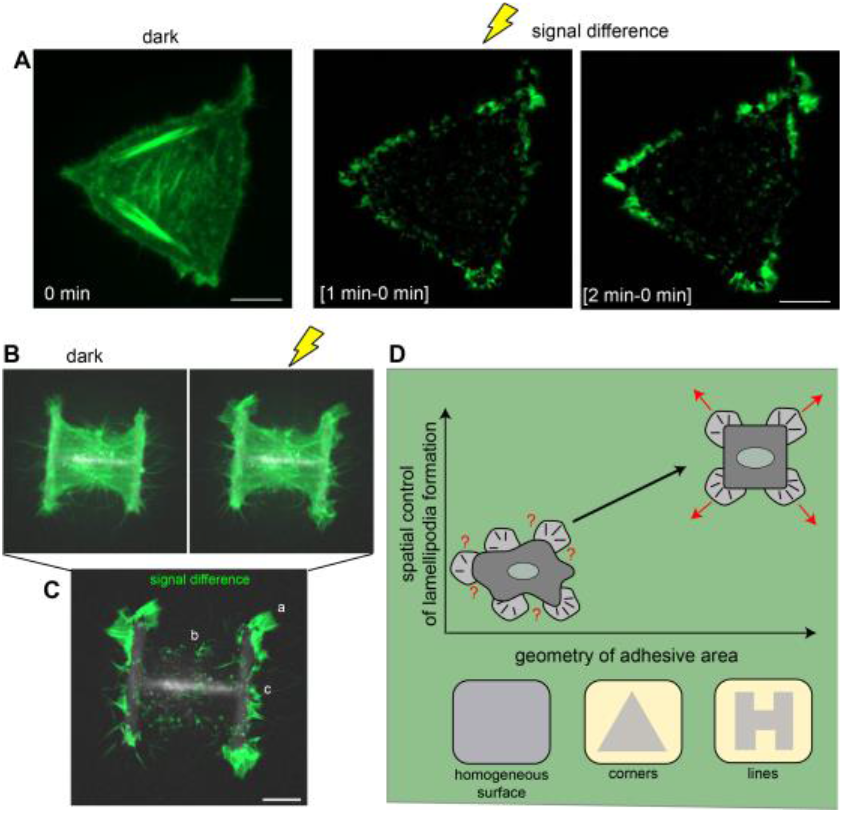
Geometry of adhesive micropatterns determine localization of light induced protrusions. (A) HFF1 cells were plated on triangular fibronectin micropatterns. Confocal images of lifeact-eGFP (green) were acquired to compute the signal difference of filamentous actin before and after photoactivation of PA-Rac1. On triangular micropatterns light-induced protrusion formation was preferentially localized to cell corners. (B) Also on H-shaped micropatterns (white) protrusion formation was localized to cell corners. (C) Computing the signal difference of lifeact-eGFP at the beginning and after 2minutes illumination demonstrates three different efficiencies of protrusion formation on H-patterns. (a) Extensive protrusions typically developed at the corners of the cell. (b) No protrusion formation was observed at non-adhesive boundaries of HFF1 cells. (c) Moderate protrusion formation was observed at fibronectin attached boundaries of HFF1 cells. (D) Schematic model for how spatial control of light-induced lamellipodia depends on the geometry of adhesion patterns. On homogeneous substrates the number and location of light-induced protrusions is not regulated, while micropatterns predefine where protrusions are formed upon PA-Rac1 activation with light. (A-C) Scale bars: 10 μm

In addition to constraining adhesion geometry, we also manipulated the distribution of cytoskeletal networks in a cell using patterns which mediate cell spreading across areas with two distinct degrees of adhesiveness. Specifically, we used H-shaped patterns on which cells spread into squares with two adhering and two non-adhering sides to characterize protrusion formation in the context of low and high adhesion (Fig. 2B). Upon photo-activation of Rac1 cells plated on these H-patterns, we observed protrusions preferentially formed at cell corners (Figure 2C). Overall, the responses to optogenetic stimulation fell into three different groups, similar to the classes of light-induced protrusion observed on homogeneous substrates. Firstly, the most active zones of protrusion were the four corners of the cell. Thereby, the spatial localization to cell corners was even more specific on H-shaped micropatterns as compared to triangular patterns. Secondly, the two cell sides overlying the micro-patterned fibronectin displayed weak protrusive activity. Thirdly, the cell sides with no underlying adhesion molecules were almost completely quiescent (Fig 2B). This pattern of light-induced lamellipodial protrusion was reversible. Combining micropatterns and optogenetics is thus an intriguing tool to control the spatial response to global optogenetic activation of signaling molecules: In addition to controlling the time of protrusions formation through light, our approach regulates the number and locations of protrusions by manipulating the actin cytoskeleton through patterned adhesion (Fig 2C).

## The cellular microenvironment modulates bleb formation

In addition to complex, actin-filled protrusions such as lamellipodia, cells can also form spherical, actin-free protrusions called blebs. No significant blebbing was observed in cells plated on non-patterned surfaces, but on fibronectin micropatterns we also observed cells producing blebs. This observation is consistent with previous studies that report a small fraction of cells produce blebs on micropatterns(13). Our cells produced blebs in one of two patterns: either across their entire surface, or preferred at the corners (Fig. 3A, B). To determine whether confinement to small adhesive patterns induced membrane blebbing and to control for chemical effects associated with micro-patterning, we created large and small triangular patterns. Consistent with confinement-induced blebbing, we observed that cells on smaller fibronectin micro-patterns showed increased blebbing activity (Fig. 3C). To characterize the actin cortex dynamics underlying cells with blebs at their corners, we acquired time-lapse microscopy images of lifeact-eGFP. At the sides of triangular cells actin structures did not change significantly. At the corners of these cells pronounced actin cortex assembly was observed during bleb retraction (Fig 3D).

**Fig. 3:**
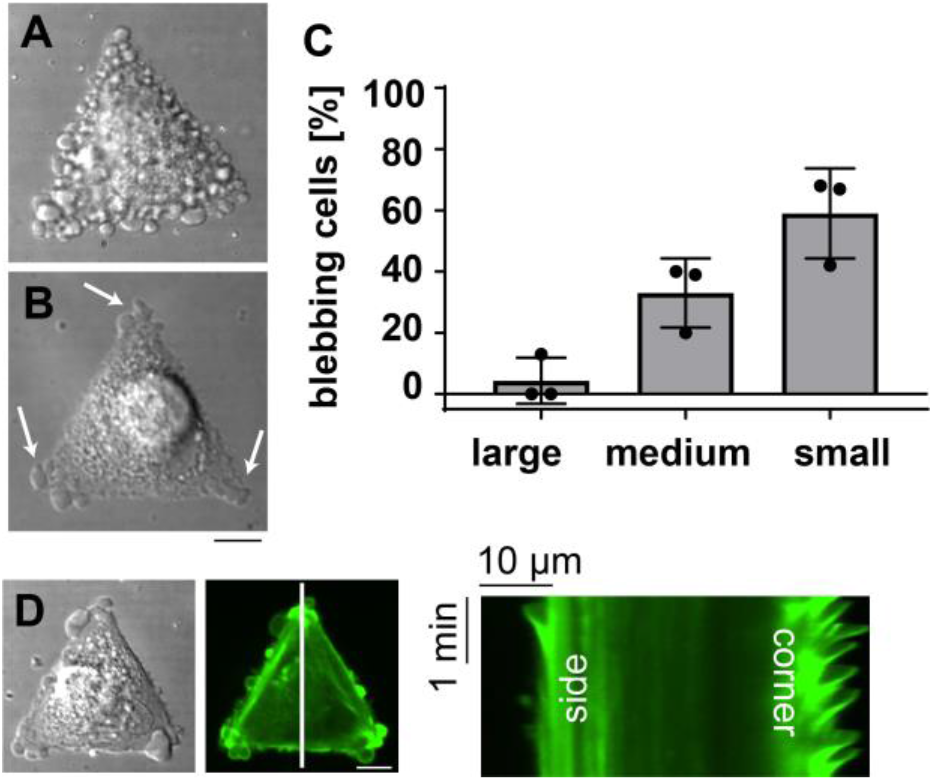
Size and geometry of micropatterns regulate bleb formation of cells. HFF1 cells were imaged 2h after seeding on triangular fibronectin patterns. Numerous cells showed blebbing activity. Thereby, blebbing covered either the whole cell (A) or was restricted to subcellular regions (B). (C) Blebbing activity depends on the micropattern area. On large micropatterns (2200 μm^^^2) cells spread typically without formation of blebs. On smaller triangles (920 μm^^^2 and 450 um^^^2) a larger fraction of cells displayed blebs. For each pattern size more than 70 cells from 3 independent samples were evaluated. The graph shows the fractions of blebbing cells in independent samples (black dots) and overall means (grey bars) with standard deviation. (D) To further characterize HFF1 cells, with blebs at subcellular region, confocal lime lapse movies of lifeact-eGFP (green) were acquired. A Kymograph computed through a corner and the opposite side of a cell (along white line)-shows intensive actin dynamics, representing bleb formation at the corner. Scale bars: 10 μm

These results demonstrate that the geometry and size of adhesive patterns modulates formation and distribution of blebs. Our findings also indicate that tissue environments with limited free space and irregular geometry of extracellular matrix proteins, may provide simple but powerful cues that dictate the morphology and locations of cellular protrusions.

## Optogenetic control of protrusion morphology on micropatterns

It has been reported, that Rac1-induced actin polymerization promotes lamellipodia formation at the expense of blebs in migrating carcinoma cells(14). We asked whether we could control the transition between blebbing and lamellipodial protrusion by optogenetically activating Rac1 in blebbing cells plated on fibronectin micro-patterns. In most PA-Rac1 cells, global activation of Rac1 converted blebs to lamellipodia within 4 minutes (Fig 4).

**Fig. 4:**
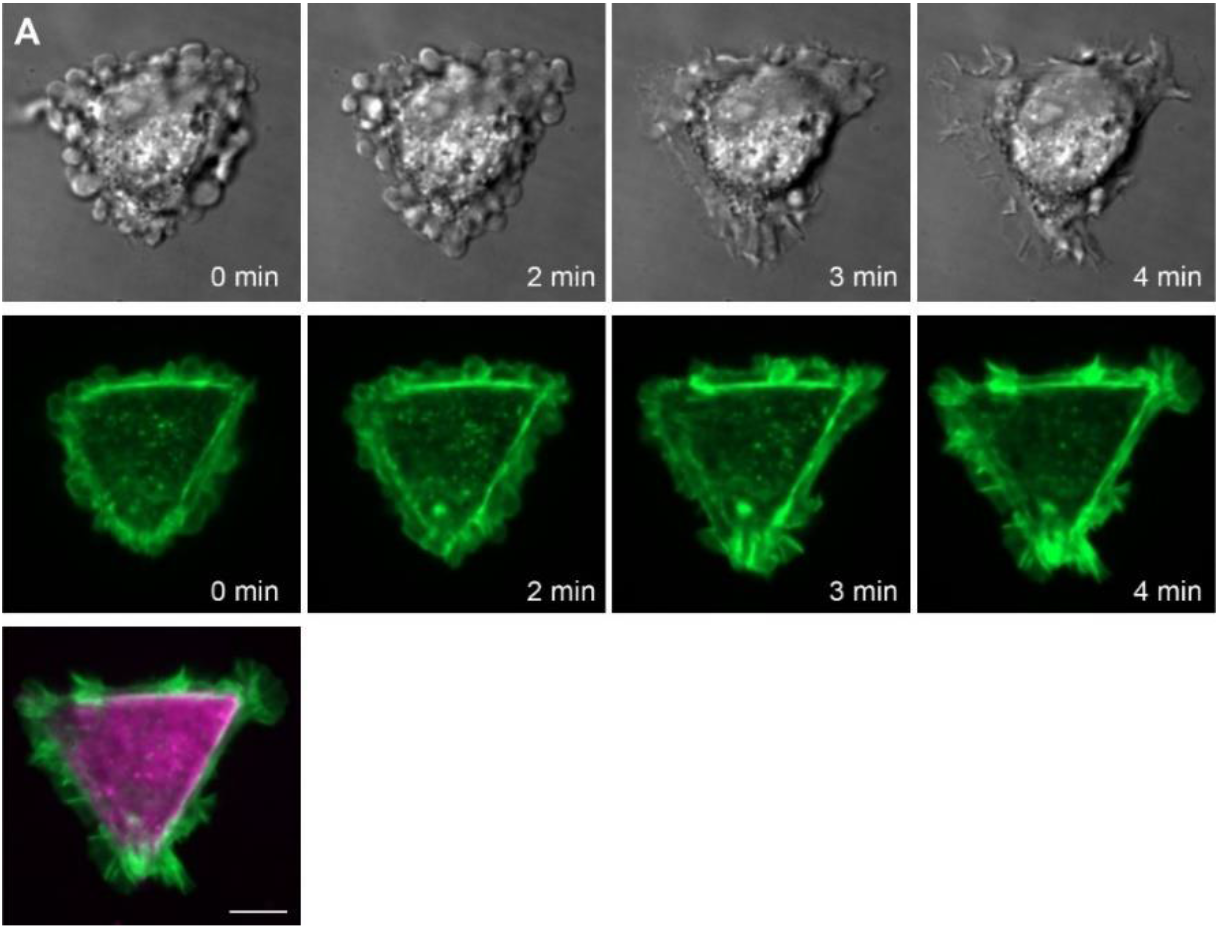
Optogenetic switching of cellular protrusion morphology. To characterize the combination of micropatterns and optogenetics as an approach for changing the morphology of blebbing cells, HFF1 cells were plated on fibronectin patterns (magenta). Time lapse images lifeact-eGFP (green) and brightfield were acquired during photoactivation of PA-Rac1. After about 3 minutes lamellipodia formed. Scale bar: 10 μm

Interestingly, however, in individual cells Rac1 activation not only failed to induce lamellipodia within 4 minutes, but actually caused bleb size to increase. This transition, in which 6 minutes of light stimulation increased the bleb volume, was reversible (Fig 5A). We used time-lapse microscopy to characterize the dynamics of these optogenetically induced “giant” blebs. In extreme cases blebs appeared to form travelling waves on the cell surface (Fig. 5B). We never observed such an increase in in bleb size in non-activated cells, which produced only small, non-traveling blebs.

**Figure 5:**
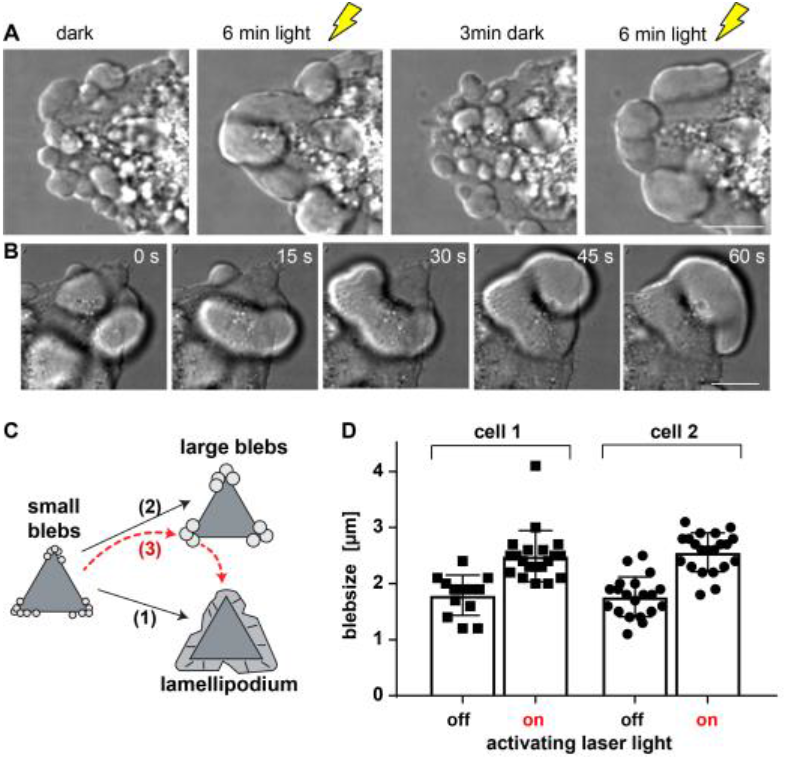
Regulation of bleb size using optogenetics. (A) Instead of forming lamellipodia upon light activation (Fig. 4), a fraction of cells formed “giant” blebs. Size modulation by photoactivation was reversible. (B) Time lapse images demonstrate that large blebs were dynamic and can travel across the cell surface. Scale bar: 10 μm (C) Schematic model for the transition of small blebs. Upon light-activation of Rac1 small blebs can be converted to lamellipodia (1) or larger blebs (2). During transitions to lamellipodia larger blebs can be observed transitionally. (D) Quantification of bleb size before light stimulation and just before lamellipodia occurred.

Notably, similar giant blebs have previously been associated with specific cell types, such as embryonic blastomere cells in which travelling blebs are commonly observed and referred to as “circus movements” (6). The origin of these circus movements and why they occur only in certain cell types has been a mystery in the field(7). Our work demonstrates that combinations of micropatterning and activation of signaling proteins provides new opportunities to custom control bleb volume and suggests, that although the size and dynamics of blebs can be vastly different among different cell types, simple biochemical modulation of protein activity can account for these dramatic differences.

An open question, however was why the two different phenotypes of larger blebs vs. lamellipodia occurred when activating PA-Rac1. To characterize the relationship of protrusion transitions from small to large blebs and from small blebs to lamellipodia, we quantified the size of blebs at cell corners that produced lamellipodia upon light stimulation. Lamellipodia formation occurred typically fast (within few minutes). Measurements were performed on such cell corners which initiated lamellipodia between 2.5 and 4 minutes. Interestingly, we observed cells in which bleb size first increased before lamellipodia formation was initiated. (Fig 5D, Supplemental figure 1). These results demonstrate that lamellipodia and an increase in bleb size can both occur within the same cell because of increased activity of signaling proteins and that large blebs are a transitory state between small blebs and lamellipodia. Moreover, they demonstrate that the combination of optogenetics and protein micropatterns is a powerful approach to regulate the transitions between small blebs, large blebs and lamellipodia (Fig 6).

**Figure 6:**
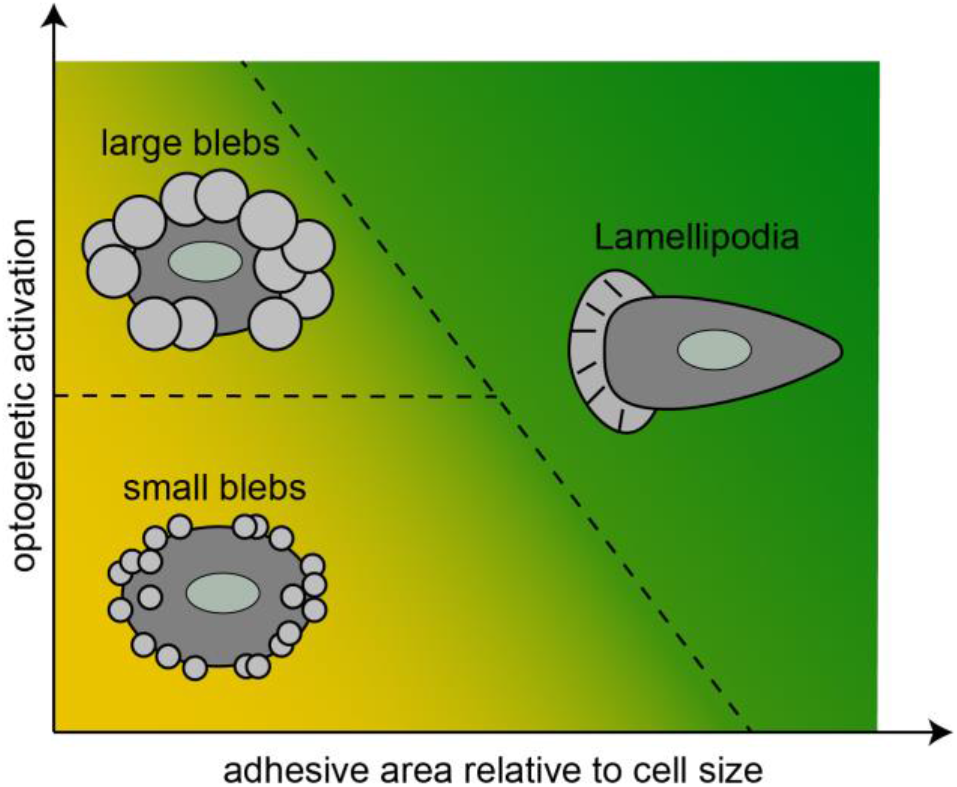
Model for how cell shape may depend on the two dimensional parameter space of PA-RacA activation and the area of adhesive patterns

## Discussion

Our results demonstrate that, when used together, optogenetics and surface micro-patterning create an effective toolkit for spatial and temporal control of cellular protrusion. Specifically, we used this toolkit to investigate three distinct morphology changes.

Firstly, we found that we could use adhesion geometry to manipulate the spatial organization of the actin cytoskeleton and thus constrain light-induced protrusions to cell corners. Previous studies demonstrated temporal and positional control of protrusion formation on homogenous substrates by positional control of laser light (10), but laser-controlled localization of protrusions is typically limited to the induction of one or few protrusions and its efficiency varies along cell boundary due to the complex organization of the cytoskeleton, as we have shown here (Fig. 1). Other studies demonstrated the capacity of micropatterns to direct protrusions to cell corners when lamellipodia formation was induced with chemicals(15), however these experiments provide limited temporal control over protrusion formation and often the exact mechanism for how chemical probes activate signaling pathways are unknown. Using a combination of whole cell illumination and micropatterns, we harvest the advantages of both techniques and demonstrate here an approach to induce arrays of protrusions with high temporal precision as well as spatial control. Notably, while the heterogeneity of optogenetic activation within cells has received little attention in the literature, our work uses the molecular complexity of cells and the resulting heterogeneous efficiency of optogenetic activation across the cell body to implement controlled protrusion localization (Fig 2). Characterizing and controlling heterogeneity of optogenetic activation may thus lead to new insights into cell function – e.g. how actin structures prime subcellular regions for future protrusion formation – and novel opportunities for technology development.

Secondly, we used optogenetics and micropatterning to control transitions between lamellipodia and blebs in fibroblasts. Optogenetic switching of cell migration has been achieved previously(14), but the integrated optogenetic and micropattern-mediated control of protrusion morphology in fibroblasts – a common model system for flat lamellipodial protrusions – represents a new level of engineering control over cell morphogenesis. We demonstrate that simple changes in the cellular environment and activation of signaling pathways can induce transitions between distinct morphologies. Notably, some cancer cells have been reported to switch between lamellipodia-driven and bleb-driven cell motility depending on environmental conditions(16)(1). Thus, our observations might not only be of relevance in the context of basic research on protrusion formation and assay development, but also for biomedical research and human health.

Finally, we demonstrate optogenetic control over bleb size. Cell type-specific variations in bleb size have been observed for decades. For example, the large blebs of blastomere cells are not only larger than ‘standard blebs’ observed in fibroblasts, but also fewer in number. These giant blebs have also been reported to circulate around the cell periphery(6). How different cell types produce blebs with different morphologies and dynamics has long been a subject of speculation. One suggestion has been that a smaller effective pore size of the cytoplasm in specific cell types could make it energetically more costly to push fluid inside blebs back into the bulk cytoplasm and thereby favor travelling blebs, while other possibilities include differences in cell cortex mechanics(7)(8). Here, we demonstrated distinct bleb sizes in the same cell through optogenetic activation, but without modulation of the cytoplasm or protein abundance. Thus, distinct pore sizes of the cytoplasm are not required to account for bleb size. Our experimental results suggests an alternative mechanism for bleb size regulation and raise an important question as to what mechanism may be responsible for differences in bleb size and dynamics. One speculation is that the abundances of distinct actin nucleators plays a role in this process: Previous proteomics and knockout experiments demonstrate that cortex assembly in blebs is mediated by the two actin nucleators mDia1 and Arp2/3(8). However, while depletion mDia1 or Arp2/3 results in increased and decreased bleb size, respectively, it is unknown whether the underlying molecular mechanism for distinct bleb size in these experiments is based on (i) the overall presence of these nucleators or (ii) their relative abundance.(8) Our results demonstrate that blebs become also larger if Rac1 – a positive regulator for branched actin synthesis by Arp2/3 – is optogenetically activated und thus has a qualitatively similar effect as the downregulation of mDia1. Thus, in contrast to two actin nucleators playing distinct roles in actin cortex synthesis, our experiments point towards a role of actin nucleator homeostasis in bleb size control. One possibility is that the distinct network architectures of the cellular actin cortex that arises in the presence of excess Arp2/3 as compared to mDiaI facilitates propagating cracks in the actin cortex.

In summary we have established the combination of micropatterns and optogenetics as a toolkit to control and interrogate transitions of cellular protrusion morphologies. Combining optogenetics and micropatterns provides control over (i) the timing of protrusion formation, (ii) the location of protrusions, (iii) their number and (iv) protrusion morphologies. Our system maintains key aspects for the induction of morphology transitions, while minimizing the complexity of morphology transitions in tissue contexts and suggests how cell morphology depends on adhesion geometry and Rac1 signalling (Fig 6).

## Acknowledgements

The authors acknowledge funding from the National Institutes of Health (R35-GM118119) and the Howard Hughes Medical Institute. KZ was supported by a research fellowship from the Deutsche Forschungsgemeinschaft (DFG). The authors also thank members of the Mullins lab for reagents, support and advice.

## Supplemental information

**Supplemental figure 1.**
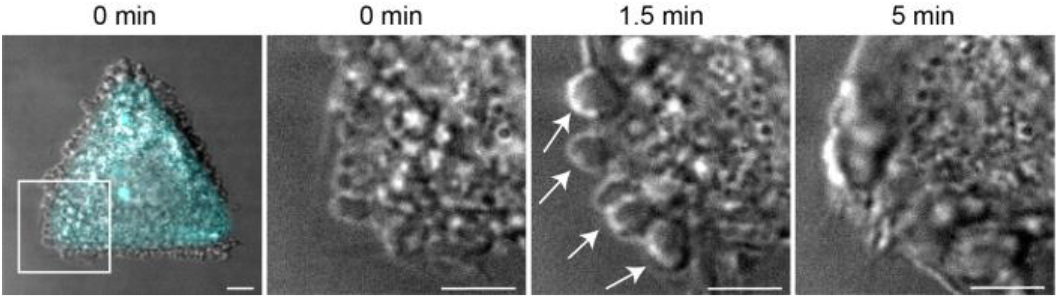
Bleb size increases during transitions from blebs to lamellipodia. Time lapse images during light-activation of PA-Rac1 demonstrate that large blebs (white arrows) represent an intermediate state between small blebs and lamellipodia. White box represents magnified region. Scale bars: 5 μm

